# Normal Reproductive function in male mice lacking pituitary kisspeptin receptor

**DOI:** 10.1101/2021.12.30.474587

**Authors:** Yaping Ma, Olubusayo Awe, Sally Radivick, Xiaofeng Yang, Sara Divall, Andrew Wolfe, Sheng Wu

## Abstract

The anterior pituitary secretion of luteinizing hormone (LH) and follicle-stimulating hormone (FSH) regulate gonadal development, gametogenesis and the secretion of the gonadal steroid hormones. The gonadotroph is primarily regulated by hypothalamic secretion of gonadotropin-releasing hormone (GnRH) from neurons of the rostral hypothalamus and is mediated by GnRH receptor signaling. Kisspeptin (KISS1)/kisspeptin receptor (KISS1R) signaling in GnRH neurons plays an essential role in reproductive function. As the kisspeptin receptor is present in the pituitary, kisspeptin signaling via the Kiss1r may regulate reproductive function at the level of pituitary. Using Cre/Lox technology, we deleted the *Kiss1r* gene in pituitary gonadotropes (PKiRKO). PKiRKO male and females have normal genital development, puberty onset, and fertility. Females have normal LH, FSH and estradiol while males had significantly increased basal serum FSH levels with no differences in basal serum LH, or testosterone levels. Overall, these findings indicate that the pituitary KISS1R does not play a role in male reproduction.

## Introduction

Pubertal onset is marked by an increase in the frequency and amplitude of gonadotropin-releasing hormone (GnRH) pulses by the hypothalamus, which is followed by increased secretion of the gonadotropins, luteinizing hormone (LH) and follicle-stimulating hormone (FSH), by the anterior pituitary (1-4). Kisspeptin and its G-protein coupled receptor (KISS1R/GPR54) have a critical role in regulating the hypothalamic-pituitary-gonadal axis in all mammalian systems (5-8). Loss of function mutations of *Kiss1r* are associated with lack of puberty onset and hypogonadotropic hypogonadism in humans and rodents, whereas gain-of-function mutations produce precocious puberty in humans (8-12). In rodents, *Kiss1r* is highly expressed in hypothalamus and its role in regulating the reproductive hormone axis is firmly established (13, 14). Conditional knock-out studies have clearly defined a role for *Kiss1r* expression in the GnRH neuron for reproductive development and fertility (15, 16). Besides the hypothalamus, pituitary and gonads, *Kiss1r* is also detected in placenta, liver, pancreas and intestine.

GnRH neurons located in the hypothalamus have long been thought to be the final neuroendocrine output regulating the reproductive axis. The pituitary incorporates diverse cues and signals to affect reproductive output including estradiol (17), testosterone (18), progesterone (19) and inhibin (20), metabolic hormones such as insulin (21, 22), and pituitary derived activin (23). Hypothalamic regulatory factors such as corticotropin-releasing hormone (CRH) (24), dopamine (25), and pituitary adenylate cyclase-activating peptide (PACAP) (26) also affect pituitary function, suggesting that the pituitary may be accessible to KISS1, to activate KISS1R signaling cascades.

Kisspeptin increases gonadotropin gene expression in mouse primary pituitary cells in culture (14). *Kiss1r* expression is enhanced in the pituitary of female mice during the estradiol-induced LH surge (27), suggesting a possible role for kisspeptin as part of the constellation of regulatory inputs to the pituitary required for the preovulatory release of gonadotropins. We sought to examine the physiological role of pituitary kisspeptin receptor using a novel kisspeptin receptor knockout mouse model (PKiRKO mouse). We find that pituitary kisspeptin signaling is not necessary for reproductive function.

## Materials and Methods

### Generation of Gonadotrope-specific Kiss1R knockout mice (PKiRKO)

To generate pituitary Kiss1 receptor (Kiss1R) knockout (PKiRKO) mice, we crossed *Kiss1r* heterozygous (fl/wt) female mice (16) with αGSU transgenic (αCre+/-) male (22, 28) mice. The αGSU transgenic (αCre+/-) effectively deletes floxed genes specifically in pituitary gonadotrophs. F1: female mice (Kiss1R fl/wt; αCre+) and male mice (Kiss1R fl/fl; αCre-) were crossed to produce PKiRKO mice (Kiss1R fl/fl; αCre+). Litter mates (*Kiss1r* fl/wt; αCre- and *Kiss1r* fl/fl; αCre-) were used as controls (referred as WT). DNA was extracted as described previously (29). Genotyping primers were designed to detect the presence of the floxed allele, WT allele, or knockout allele of Kiss1r: P1 sense (located in exon 1) and P3 antisense (located in exon 3). Genomic DNA obtained from extra-pituitary organs (eg, tail), primers P1 and P3 amplify a 2096-bp amplicon to indicate the floxed Kiss1r allele and 1882-bp band to indicate the WT allele. Genomic DNA obtained from the pituitary using primers P1 and P3 will amplify a 1120-bp amplicon if the sequence between the LoxP sites is excised indicating a KO allele.

### Animals

Adult male and female mice (>2 months old) were used in this study. All animal studies were carried out in accordance with National Institutes of Health guidelines on animal care regulations and were approved by the Animal Care and Use Committee of the Johns Hopkins University. Mice were maintained under constant conditions of light and temperature (14: 10 h light/dark cycle; 22 C) and were fed a normal chow and water ad libitum.

### Quantitative Real-Time PCR (qPCR)

RNA was extracted from the pituitary, liver, ovaries, uterus, testes, epididymis, adipose tissue and from two hypothalamic fragments encompassing the arcuate and the anteroventral periventricular nucleus (AVPV) (30), using TRIzol reagent (Ambion life technologies, Carlsbad CA, USA), according to the protocol provided by the supplier. 1µg of RNA was reverse transcribed to cDNA using an iScript cDNA kit (Bio-Rad Laboratories). Real-time qPCR was performed to determine the presence and relative expression levels of *Kiss1*r mRNA in the various tissues. Real-time qPCR was performed in duplicate using SYBR Green Master Mix (Bio-Rad Laboratories) and the CFX Connect qPCR machine (Bio-Rad Laboratories). For each primer set, PCR efficiency was determined by measuring a 10-fold serial dilutions of cDNA and reactions having 95% and 105% PCR efficiency were included in subsequent analyses. Relative differences in cDNA concentration between WT and PKiRKO mice were then calculated using the comparative threshold cycle (C_t_) method. To compare the difference of *Kiss1*r expression in the same tissue between WT and PKiRKO, a ΔC_t_ was calculated to normalize for internal control using the equation: C_t_ (Kiss1R) – C_t_ (18S). ΔΔC_t_ was calculated: ΔC_t_ (PKiRKO) -ΔC_t_ (WT). Relative Kiss1R mRNA levels were then calculated using the equation fold difference = 2^ΔΔ^C_t_.

### Pubertal onset and assessment

Preputial separation (PPS) in males was assessed daily beginning after postnatal day 21 by determining whether the prepuce could be manually retracted with gentle pressure. PPS is testosterone dependent and thus is an indicator of activation of the reproductive axis in males. Puberty in rodents is dependent on weight (31); hence, the weights of PKiRKO and wild type were assessed once a week in prepuberty (day 21) through adulthood (day 49). Anogenital distance is testosterone dependent and was determined at 8 weeks of age. Wet testicular weights were determined in freshly dissected mice at two months of age. In females, age at vaginal opening and first estrus are two markers of puberty onset and were assessed for daily in mice after 21 days of life until achieved. Vaginal smears were obtained daily over a period of 14 consecutive days in 6- to 11-week-old mice and cellular morphology examined under microscope.

### Hormone assays

Blood samples were collected from submandibular vein (28) between 9:00 and 10:00 AM and basal levels of serum LH and FSH were measured. LH and FSH were measured using a Milliplex MAP immunoassay (Mouse Pituitary panel; Millipore, Massachusetts) on a Luminex 200IS platform (Luminex Corporation). The assay detection limit for LH was 0.012 ng/mL and for FSH was 0.061ng/mL. The intra-assay coefficients of variation (CV) for LH and FSH were between 5 % and 9%. Serum E_2_ levels were measured with a mouse/rat E_2_ kit from Calbiotech (El Cajon, CA). The sensitivity for E_2_ was 3 pg/mL. Testosterone levels were measured by radioimmunoassay by the University of Virginia Ligand Assay Core (Charlottesville, Virginia).

### Fertility assessment

To determine whether male PKiRKO mice were fertile relative to controls, 3 WT and 4 PKiRKO male mice were housed individually with proven fertile WT female mice for 14 consecutive days and then were separated. If a pregnancy ensued, one week after female gave birth, a new male was inserted. Males were rotated among the proven females. To assess fertility in females, a similar strategy was used. Females were housed individually with proven fertile WT male mice for 14 consecutive days and then separated. If a pregnancy ensued, one week after female gave birth, a new male was inserted, rotating among the 7 proven males. The duration of the fertility study was 4 months with 4 rotations for each mouse.

## Data Analysis

Data were analyzed by two tailed unpaired student t tests using Prism software (GraphPad Software, Inc, La Jolla, CA). All results are expressed as mean ± SEM (standard error of the mean). P < 0.05 was defined as statistically significant.

## Results

### Generation of pituitary specific *Kiss1r* knockout mice

PKiRKO (Kiss1R^fl/fl^; αCre^+^) mice were generated by Cre recombinase mediated excision of exon 2 of the *Kiss1r*, resulting in an attenuated *Kiss1r* gene in pituitary, as shown in the schematic diagram of Figure 1A. The PCR product indicates the homozygous floxed-*Kiss1r* alleles (2096 bp) and WT alleles (1882 bp) (Figure 1 B); both bands are present in the heterozygous floxed-*Kiss1r* mouse schematized by Novaira HJ *et al*. (16). Shown in Figure C Lane 1 is a WT αCre-male mice, followed by two αCre+ male mice in lanes 2 and 3.

**Figure 1.**
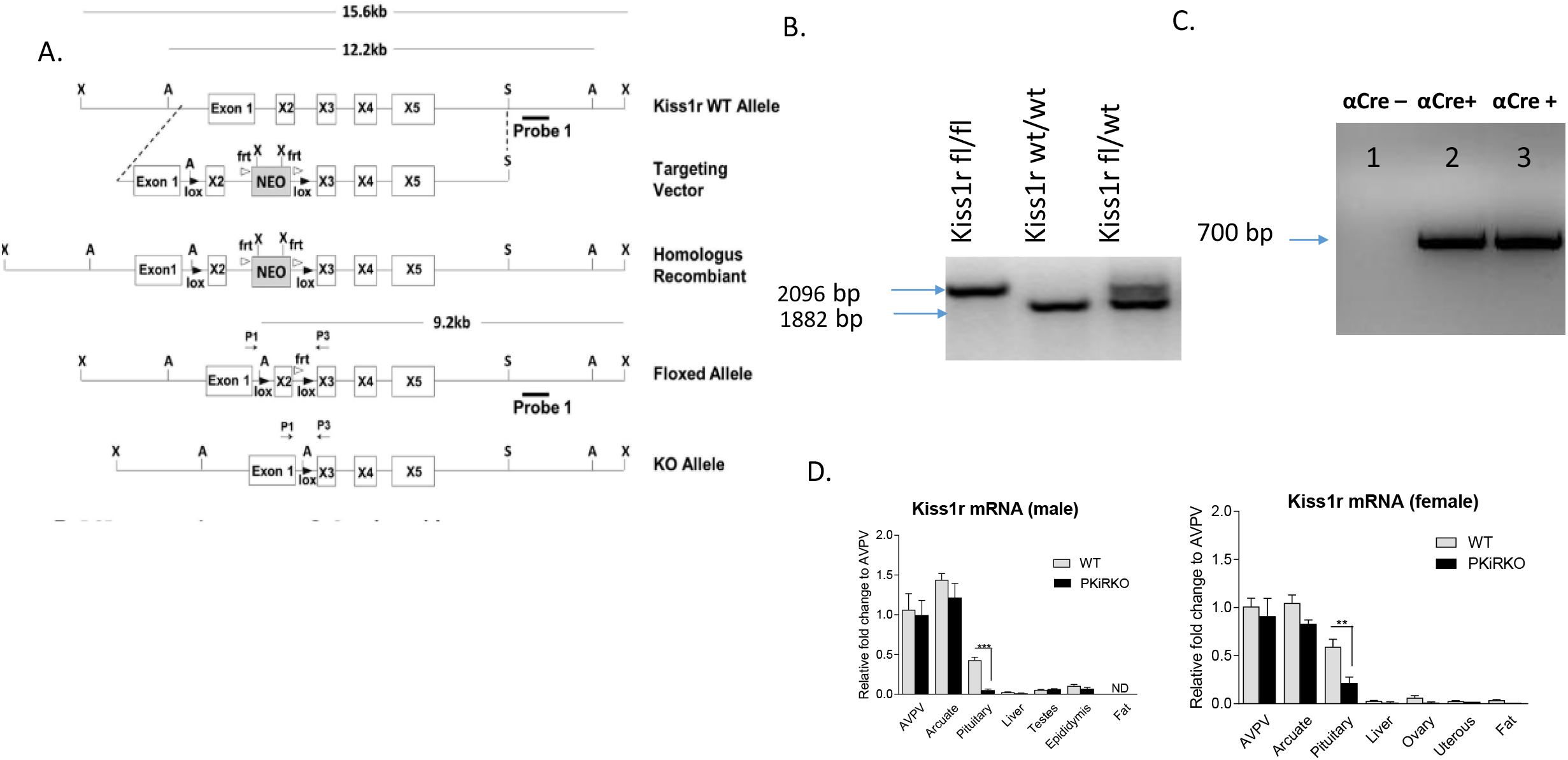
Development of the PKiRKO mouse. A. schematic diagrams of constructs used to generate PKiRKO mice. Mice bearing LoxP sites flanking exon 2 of the Kiss1r were crossed with transgenic mice expressing Cre recombinase specifically in Gonadotrophs (αGSU). B. Genotyping by PCR analysis of the genomic DNA produced a 2096 bp amplicon in mice with a floxed allele and an 1882 bp amplicon in kisspeptin fl/fl mice, also shown are both bands present in the heterozygous floxed-*Kiss1r*. C. Lane 1-αCre^+^ transgene negative. Lane 2 & 3-αCre^+^ transgene positive mice. Amplicon size is 700 bp. D. Quantitative RT-PCR analysis of Kiss1r mRNA extracted from male and female mouse tissues. *Kiss1r* was significantly reduced (87.6% for male, 63.7% for female) in the pituitary of PKiRKO (KO) mice compared with that in wild type (WT) mice, but no difference in *Kiss1r* expression was observed in other tissues. Data are means±SEM (n=8 for pituitary, n=4 for other tissues).

### Pituitary and tissue *Kiss1r* mRNA expression in PKiRKO mice

The levels of *Kiss1r* mRNA and or protein expression was assessed by qPCR and immunostaining. qPCR demonstrated a reduction in *Kiss1r* mRNA by 88% (Figure 1D) and 64% (Figure 1E) in the pituitary of male and female PKiRKO mice (n=8), respectively, compared with WT mice (n=8). In contrast, no change in *Kiss1r* mRNA was observed in other tissues including the hypothalamus, adipose tissue, liver, or gonads (P> 0.05, n =4) (Figure 1 D&E).

### Pubertal onset and reproductive function are normal in PKiRKO mice

Body weight has been demonstrated to be highly associated with age of puberty (32). Body weight was not different between male or female PKiRKO and WT mice (figure 2A). In males, PPS (preputial separation) is an androgen dependent process that serves as an external sign of male puberty onset (31). No significant difference was observed age of PPS between the WT and PKiRKO groups (Figure 2B), or achieved anogenital distance (figure 2C). The age of vaginal opening and first estrus was not different between PKiRKO and WT groups (Figure 2D). In sexually mature mice, gonad weight is a marker of reproductive tissue function. Gonad weights were normal between PKiRKO and WT male and female mice (Figure 3 A and B). Estrous cycling pattern was similar between PKiRKO and WT mice (Figure 3C); percent time spent at each cycle stage was not significantly different. Testes or ovary histology was not different between PKiRKO and WT mice (data not shown). The number of sperm in the seminiferous tubules was not statistically different between WT and PKiRKO mice (data not shown).

**Figure 2.**
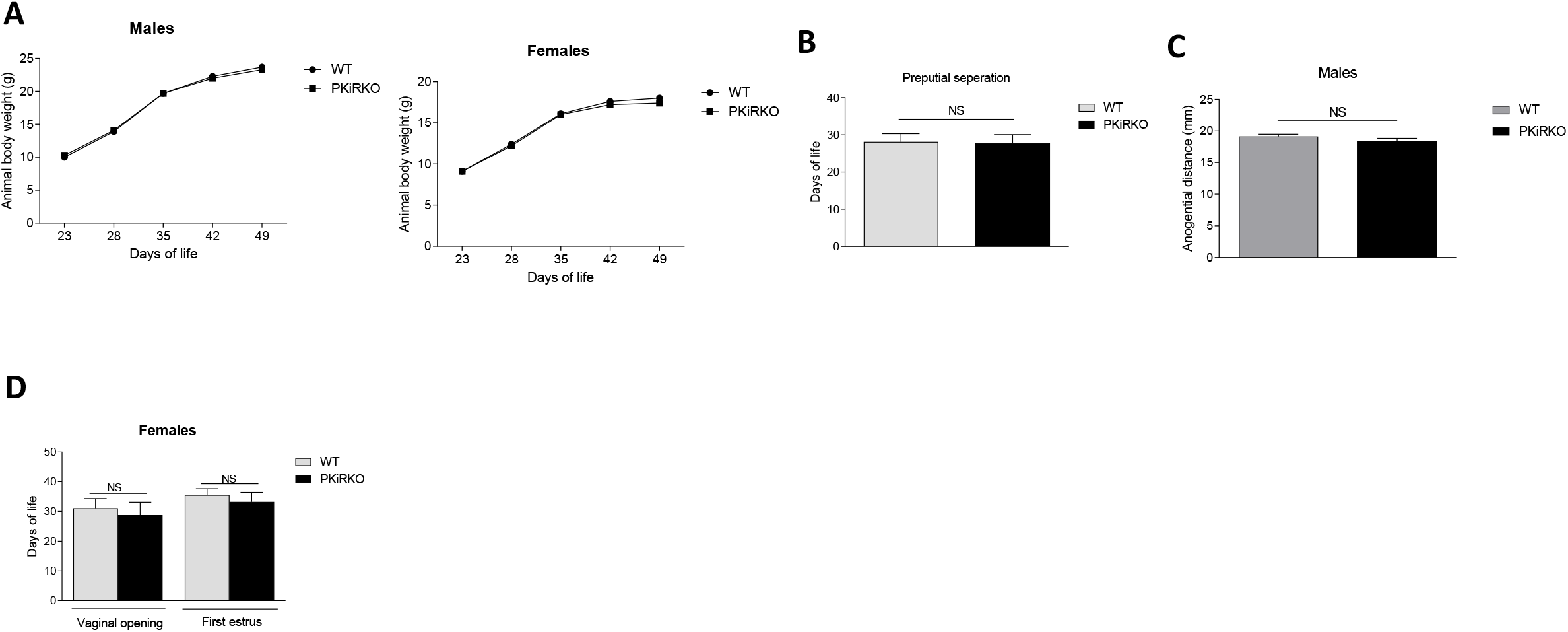
PKiIRKO mice have normal puberty onset. A.Body weight change over time was not different between PKiRKO (n=15) and WT (n=15) male mice or female PKiRKO (n=15) and WT (n=15) mice. B. No significant difference was observed in age of preputial separation (PPS) in males. (28.1 ± 2.2 [WT] vs 27.8 ±2.3 [PKiRKO]) WT (n=15) PKiRKO (n=15). C. Anogenital distance was not different between WT and PKiRKO male mice (n=6 each group). D. No difference was observed in age at vaginal opening (30.2± 2.6 [WT] vs. 28.9±3.9[PKiRKO]) or first estrus (35.6±2.4 [WT] vs 33.3±3.2 [PKiRKO]).

**Figure 3.**
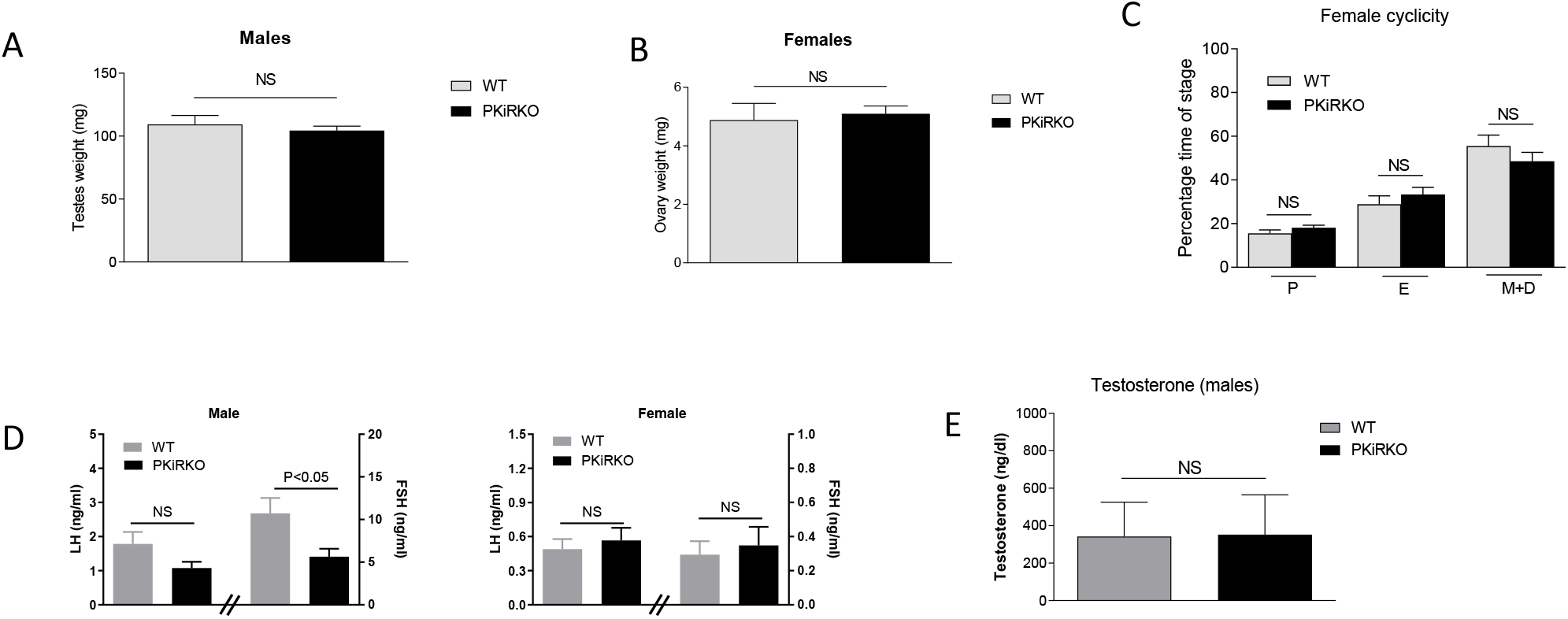
PKiRKO mice have normal reproductive function. A. Testes weight was not significantly different between PKiRKO and WT male mice (n=6). B. Ovary weight was not significantly different between PKiRKO and WT female mice (n=6) C. Vaginal cytology was examned daily over a period of 14 consecutive days in 6- to 11-week-old mice. WT and PKiRKO groups both exhibited regular estrous cycles the percentage time spent at each cycle stage did not differ between groups. N= 6 each group. D. Morning LH (left Y axis and FSH (right Y axis) in male (left panel) and female (right panel) mice. n=6, P>0.05) E.Testosterone levels are not different between PKiRKO and WT mice (n=6 each group). NS= No significant difference.

Morning serum samples were obtained from non-breeding, post pubertal mice and serum LH and FSH hormone levels were determined. Basal serum LH in PKiRKO males or females was not significantly different relative to WT mice while FSH was significantly lower in PKiRKO males (Figure 3D). Testosterone and estradiol were not significantly different between WT and PKiRKO mice (Figure 3E and F).

### Normal fertility in PKiRKO mice

Fertility was examined in WT and PKiRKO mice using a rotating mating protocol. PKiRKO male and female mice demonstrated a normal ability to produce offspring, as shown in Figure 4 A-C. Female WT mice bore their first litter with a similar latency after introduction to PKiRKO males or WT males [22.0±0.8d, *n=4* (PKiRKO) vs 22.3±0.6d, *n=* 3(WT); *P>*0.05] (Fig. 4A). Female WT mice had the same number of litters with PKiRKO and WT sires [3.8±0.5, *n=4* (PKiRKO) vs 3.7±0.6, *n=3* (WT); *P>*0.05] (Fig. 4B) and a similar number of pups per litter [(7.4±1.4, *n=4* (PKiRKO) vs 9.3±1.1, *n=3* (WT); *P>*0.05] (Fig. 4C). The same pattern was shown in males.

**Figure 4.**
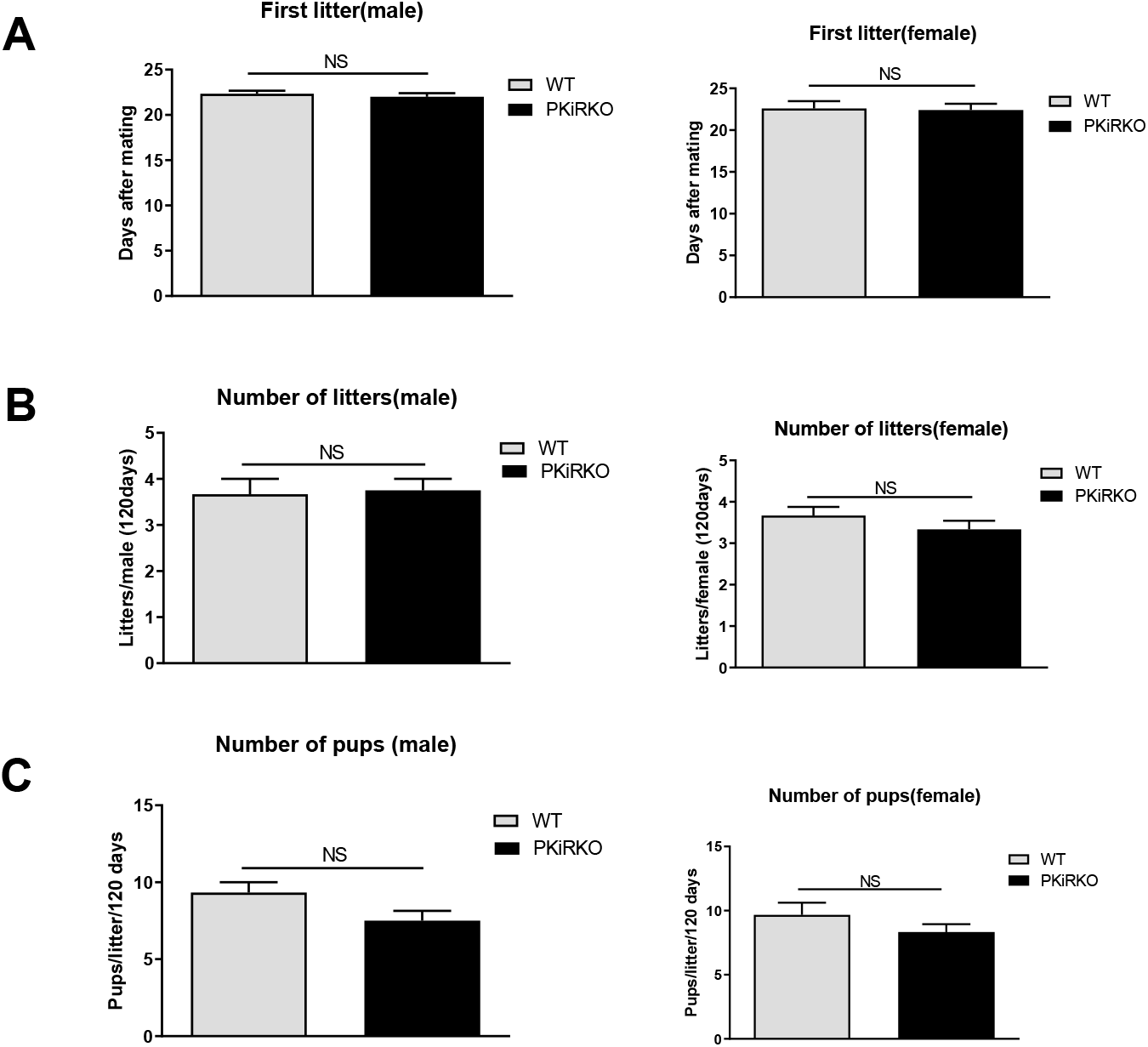
PKiRKO Mice Have Normal Fertility. A. After introduction with WT female (left panel) or male (right panel) respectively, the day of first litter was recorded. Values are mean ± S.E.M. NS, no significant. B. Total numbers of litters per male (left panel) or female (right panel) was not significantly different between WT and PKiRKO miceC. Number of pups per litter was also not significantly different between WT and PKiRKO mice (male, left panel; female right panel).

## Discussion

Kisspeptin is a potent activator of the GnRH neuron, and thus is a critical regulator of hypothalamic control of reproduction. The role of the KISS1/KISS1R signaling system has widened as investigators have identified extra-hypothalamic roles in tissues such as the liver, uterus, testes and ovary (33). Expression of KISS1R in the pituitary gonadotroph (34) points to the pituitary as a possible target for kisspeptin action. The functional significance of pituitary KISS1R is not well defined and may be the target of either centrally or peripherally derived KISS1. We sought to assess the role of KISS1 signaling at the level of the pituitary in mice by the development of a pituitary *Kiss1* knockout model (PKiRKO). We demonstrate that near complete reduction of pituitary KISSR gene and protein expression in does not affect reproductive development or function.

KISS1R is expressed on mouse pituitary gonadotrophs (FIG 2) which supports studies in rat (43) and sheep, (44) demonstrating co-expression of LH and KISS1R in the pituitary. Investigators have observed KISS1 induction of LH secretion in cultured primary rat and primate pituitary cells (35,36) and up-regulation of gonadotropin gene β-subunits, LHβ and FSHβ gene expression in LβT2 cells (14), although others have seen no direct effect of KISS1 on LH or FSH secretion (37,38). Other studies suggest that KISS1 may act directly on pituitary gonadotropes to stimulate LH release (39-41) or to increase gonadotropin or *Gnrhr* gene expression (42) which were observed *in vitro* using cell lines or primary culture. Our data indicate that these pituitary kisspeptin receptor signaling pathways may not play a role in reproduction. The PKiRKO male mice had a significantly lower morning FSH level (Figure 3). This lower FSH did not seem to affect reproductive function, as the PKiRKO mice also had normal testosterone, testes cell structure and gross sperm production (Figure 3 and data not shown). We achieved 88% reduction in kisspeptin gene expression in the pituitaries of male mice and 64% in female mice so it is possible that the reproductive phenotype of the PKiRKO mice would be more pronounced if a greater reduction was achieved. The gonadotroph comprises a minority of the pituitary cell population so kisspeptin receptor expression in other pituitary cells may account for the observed expression. The αGSU transgenic Cre mouse effectively deletes floxed genes specifically in pituitary gonadotrophs indicating that the observed kisspeptin receptor in the pituitary is likely not present in the gonadotroph.

The LH and FSH response to peripheral and central administration of kisspeptin is used to assess GnRH neuron activity. The LH and FSH of humans and mice with disorders of GnRH neuron causing hypogonadism do not rise after peripheral kisspeptin administration. These data agree with our results that there does not appear to be a direct role of kisspeptin on pituitary regulation of LH, FSH, and reproductive function; kisspeptin does not directly affect pituitary function and LH/FSH physiology in males. Kisspeptin may become available to probe GnRH neuron activity in humans to differentiate between delayed puberty and hypogonadotropic hypogonadism. Kisspeptin action is specifically on the GnRH neuron indicating that this test will be specific to probe for GnRH neuron function.

Overall this study suggests that absence of kisspeptin signaling in pituitary gonadotrophs does not affect reproductive function or fertility in male or female mice. The functional significance of kisspeptin receptor expression in the pituitary remains unclear.

